# Persistent virulent phages exist in bacterial isolates

**DOI:** 10.1101/2024.12.31.630880

**Authors:** Peter Erdmann Dougherty, Charles Bernard, Alexander Byth Carstens, Kim Stanford, Tim A. McAllister, Eduardo P. C. Rocha, Lars Hestbjerg Hansen

**Affiliations:** Department of Plant and Environmental Science, University of Copenhagen, Frederiksberg, Denmark; Institut Pasteur, Université Paris Cité, CNRS UMR3525, Microbial Evolutionary Genomics, Paris, 75015, France; Department of Biology, University of Lethbridge, Lethbridge, Alberta, Canada; Agriculture and Agri-Food Canada, Lethbridge Research and Development Centre, Lethbridge, Alberta, Canada T1J4B1

## Abstract

Despite the immense diversity of tailed bacteriophages, they are traditionally classified as either virulent or temperate, with only the latter thought capable of long-term persistence in bacterial cells through lysogeny. Virulent phages, characterized by their obligatory lytic cycle, are assumed to lack the ability to persist within bacterial colonies, and their infection is expected to decimate the host population under *in-vitro* conditions. Consequently, when bacterial isolates are cultured for sequencing, the resulting assemblies are not expected to contain virulent phage genomes. To test this assumption on a large scale, we analyzed over 267,000 publicly available *Escherichia* assemblies. Surprisingly, we identified 373 genomes corresponding to virulent phages within the bacterial genomes. These viral genomes are associated with specific phage groups and especially with jumbo phages with very large genomes (>200 kb). Chimallin was a core gene in two of these jumbo phage clusters, the major protein used by some jumbo phages to form a protective phage nucleus during infection. We found multiple lines of evidence suggesting that these virulent phage genomes in bacterial assemblies arise from persistent infections rather than contamination. Supporting this, we experimentally demonstrated the coexistence of non-temperate jumbo phages with their bacterial hosts. In a targeted follow-up search for three clades of persistent jumbo phages, we identified 285 additional jumbo phage genomes in bacterial taxa beyond *Escherichia,* highlighting that there are many more undiscovered persistent phages in bacterial assemblies. Our findings challenge the traditional virulent-temperate dichotomy, highlighting the overlooked diversity and prevalence of non-canonical phage lifestyles.

## Introduction

Bacteriophages, the viruses of bacteria, are the most diverse and abundant entities on Earth^1,2^. A key factor contributing to this ubiquity is the flexible relationship in which phages engage their hosts, ranging from antagonistic to conditionally mutualistic^3^. Phages are generally categorized into three lifestyles: virulent, temperate, and chronic^4^. Virulent phages (sometimes called lytic phages) replicate exclusively through the lytic cycle, entering the bacterial cell to immediately produce new viral copies and subsequently lyse the host cell. Temperate phages also lyse host cells but may first enter the lysogenic cycle, lying dormant as prophages either integrated within bacterial chromosomes or maintained as plasmids^4^. Prophages may carry genes beneficial to their bacterial hosts such as virulence factors^5^ or anti-phage defense systems^6^, and may even facilitate warfare between bacterial strains^7^. Lastly, filamentous phages of the Inoviridae family display a chronic lifestyle, with virion continuously released from the infected host cell without cell death^8^.

Not all phages fit neatly into this orderly system. By the 1940s it was already clear that the virulent Escherichia phage T4 delays lysis of its host cell in response to high multiplicity of infection (MOI), albeit without host cell division^9,10^. Later, a key T4 lysis inhibition gene, *rI,* was associated with long-term co-existence in slow-growing cells^11^. Similarly, *Pseudomonas* phiKZ-like phages appear to also be capable of MOI-dependent lysis inhibition in growing cells^12,13^. In a different example, the temperate Salmonella phage P22 can form temporary episomes that are asymmetrically inherited upon cell division, giving rise to a temporarily immune, uninfected bacterial sub-population which results in the maintenance of a non-infected bacterial host population^14^. Employing a different mechanism, the virulent *Bacillus* phage phi29 postpones its lytic cycle in response to the host sporulation regulator protein Spo0A, thus avoiding reproduction in environments unfavorable to both host and phage^15^. The most abundant phage family in the human virome, the *Bacteroides* crAssphages^16^, also have unusual lifestyles with long-term coexistence between host and phage. Rapid phase variation in host capsular polysaccharides has been shown to contribute to coexistence between *Bacteroides intestinalis* and CrAss-like phage ΦCrAss001 by maintaining susceptible and resistant subpopulations^17^. Recently, a crAssphage isolated through a plaque-free protocol was also shown to display a more temperate nature, maintained episomally as a phage plasmid without causing major cell lysis^18^.

Categorizing these diverse, alternative lifestyles of bacteriophages is challenging. The terms “pseudolysogeny” and “carrier state” have variously been used to describe such alternative phage-host interactions. “Carrier state” has generally been used to describe specific cases such as P22 or ΦCrAss001, where population-level coexistence of phage and bacteria is mediated by subpopulations of resistant and susceptible cells^19–21^. In contrast, “pseudolysogeny” lacks a consistent and has been used somewhat as a catch-all for poorly understood alternative phage-host interactions^19,20,22^. Due to these inconsistencies and the lack of a consensus umbrella term for the examples described above, we use the term **persistent infection** to describe a non-lysogenic infection that persists within the host population. This term refers to a phage-host interaction rather than the phage itself and is unrelated to the chronic infections of filamentous phages.

Practical limitations are a major impediment to the study of persistent infections. Most lab protocols are designed around the plaque assay, which works well for phages that quickly clear bacterial populations and produce high viral titres. In contrast, phages that persist within their hosts may not form plaques^17^ and we therefore have little idea of how common persistent infections are beyond individual case studies. Here, we hypothesize that persistent infections may remain undetected during routine bacterial isolation, culturing, and sequencing due to coexistence between the phage and host. Accordingly, the presence of non-temperate phage genomes in bacterial assemblies could be indicative of persistent infections. We were inspired to test this hypothesis during our recent isolation and sequencing of an *Erwinia* jumbo phage^23^ that formed tiny, faint plaques. When searching public databases for genomes similar to the jumbo phage, we were surprised to see hits from bacterial assemblies. To investigate this systematically, we conducted a search for non-temperate phage genomes in 267,092 *Escherichia* genome assemblies from GenBank. Our search uncovered hundreds of previously undetected, high-confidence virulent phage genomes in bacterial assemblies, with multiple lines of evidence supporting their interpretation as persistent infections. These results emphasize the importance of considering phage-host interactions as a gradient of outcomes instead of the binary labels of lysis or lysogeny.

## Results

### Non-temperate bacteriophage genomes are found in *Escherichia* genome assemblies

We searched 267,902 GenBank assemblies from *Escherichia* isolates for virulent phage contigs. *Escherichia* was chosen for two reasons; it is both the second-most sequenced bacterial genus (after *Salmonella*) and the second-most frequent phage isolation host (after *Mycobacterium*) and thus provided large datasets of both phage and bacterial genomes. Within these bacterial assemblies, we identified 28,819 contigs with >25 proteins sharing at least 70% of their proteins with the 4,890 sequenced phage genomes from GenBank and classified by BACPHLIP^24^ as virulent. The contigs were then filtered multiple times to ensure they represented virulent phages. First, we discarded contigs that were more similar to known temperate phage genomes than to known virulent phage genomes. Next, we labelled contigs that may resemble un-sequenced temperate phages, using Sourmash^25^ to identify 4,726 contigs similar to regions within complete *Escherichia* genomes in the GenBank database.

We then used vCONTACT2^26^ to build a protein-sharing network of the contigs with all sequenced *Escherichia* phage genomes. We identified large clusters within this network using Leiden clustering^27^. Contigs within the same Leiden cluster and from the same bacterial assembly were binned as single phage genomes, as these likely represent incomplete phage genomes. Clusters containing only temperate phage isolates were discarded, while those with both temperate and virulent isolates were manually filtered to ensure only clusters of high-confidence virulent phages remained. This process resulted in a trimmed network of 1,199 virulent phage isolates were clustered with 373 non-temperate phage genomes from bacterial assemblies (Figure 1A). The 373 non-temperate phages (Supplementary table S1) consisted of 493 contigs from 370 bacterial assemblies.

**Figure 1.**
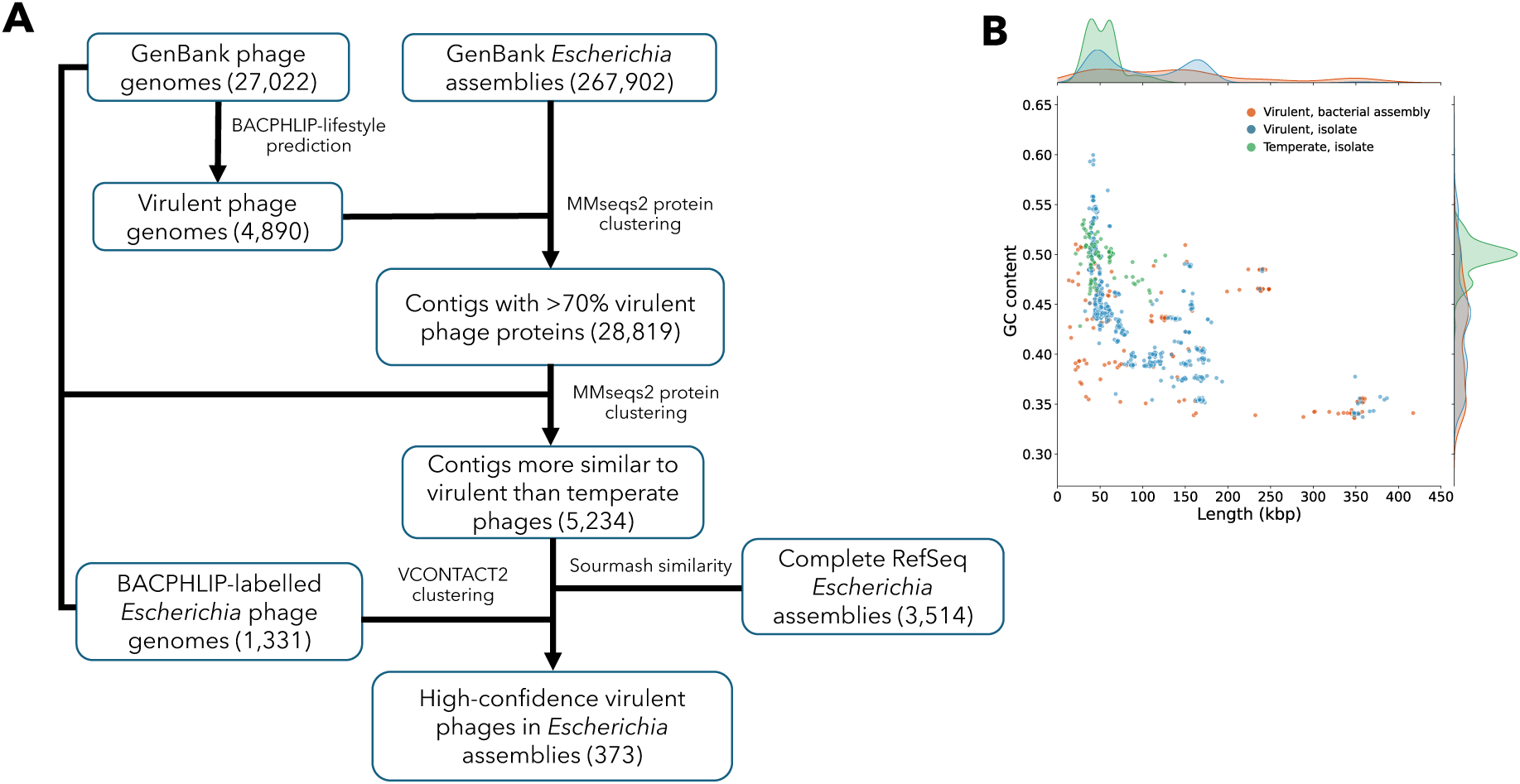
**A** Overview of the workflow used to identify virulent phage contigs in *Escherichia* isolate assemblies from GenBank. **B** Scatterplot of genome length vs GC content for three groups of phages; persistent phages found in bacterial assemblies (N=273), virulent phage isolates (N=1199), and temperate phage isolates (N=120). Relative distributions for each of the three groups also shown.

We conducted additional quality-control checks, first checking for bacterial contamination by estimating the number of bacterial genes in each contig using checkV^28^. The number of bacterial genes in the 373 non-temperate phages ranged from 0 to 3 with a mean of 0.2, indicating very little bacterial contamination. We also analyzed the average GC content and length of the non-temperate phages (Fig. 1B), as temperate *Escherichia* phage genomes from isolated phages generally exhibited higher GC content (mean 0.50±0.01) compared to virulent *Escherichia* phage isolates (0.43±0.06). The average GC content of the 373 non-temperate phages from bacterial assemblies was 0.42±0.05, consistent with the hypothesis that these are virulent phages. The temperate phage isolates were also on average shorter than the virulent phage isolates (54±20 kbp and 101±61 kbp respectively), while the 373 non-temperate phages were even longer (142±99 kbp).

Finally, we ensured these contigs were not phage-plasmids by searching for hits of HMM^29^ profiles for 38 plasmid replication proteins and 9 partition system proteins as previously described^30^. We found only 15 hits to ParB and one hit to the Rop regulator, both protein families that contain a promiscuous HTH DNA binding domain. In each case, the protein hits also lacked their cognate components (ParA and XYZ, respectively). All these tests support the classification of the 373 phages as non-temperate. Given the strictness of our filtering regime, it is likely that this number is an underestimate, and that there may be many more undiscovered non-temperate phages in bacterial genomes.

### Non-temperate phage genomes within Escherichia assemblies correspond to persistent infections rather than contamination

Having shown the phages are not prophages or phage plasmids, we next assessed the possibility that they might represent lab contaminations rather than persistent infections originating prior to bacterial isolation. First, we analyzed the ratio of viral read coverage to bacterial read coverage in the 232 bacterial assemblies for which reads were available (Supplementary table S1). We reasoned that if the ratio of persistent phage to bacterial genome coverage was much smaller than 1, the phages might represent non-replicating contaminants. Conversely, a ratio much greater than 1 could highlight phages undergoing canonically virulent lifecycles (Fig. 2A). Reassuringly, the median relative ratio was 2.8, with only one phage below 0.1 and none above 100. These results are consistent with the hypothesis that these are neither contaminants nor classically virulent phages actively replicating at the population scale.

**Figure 2.**
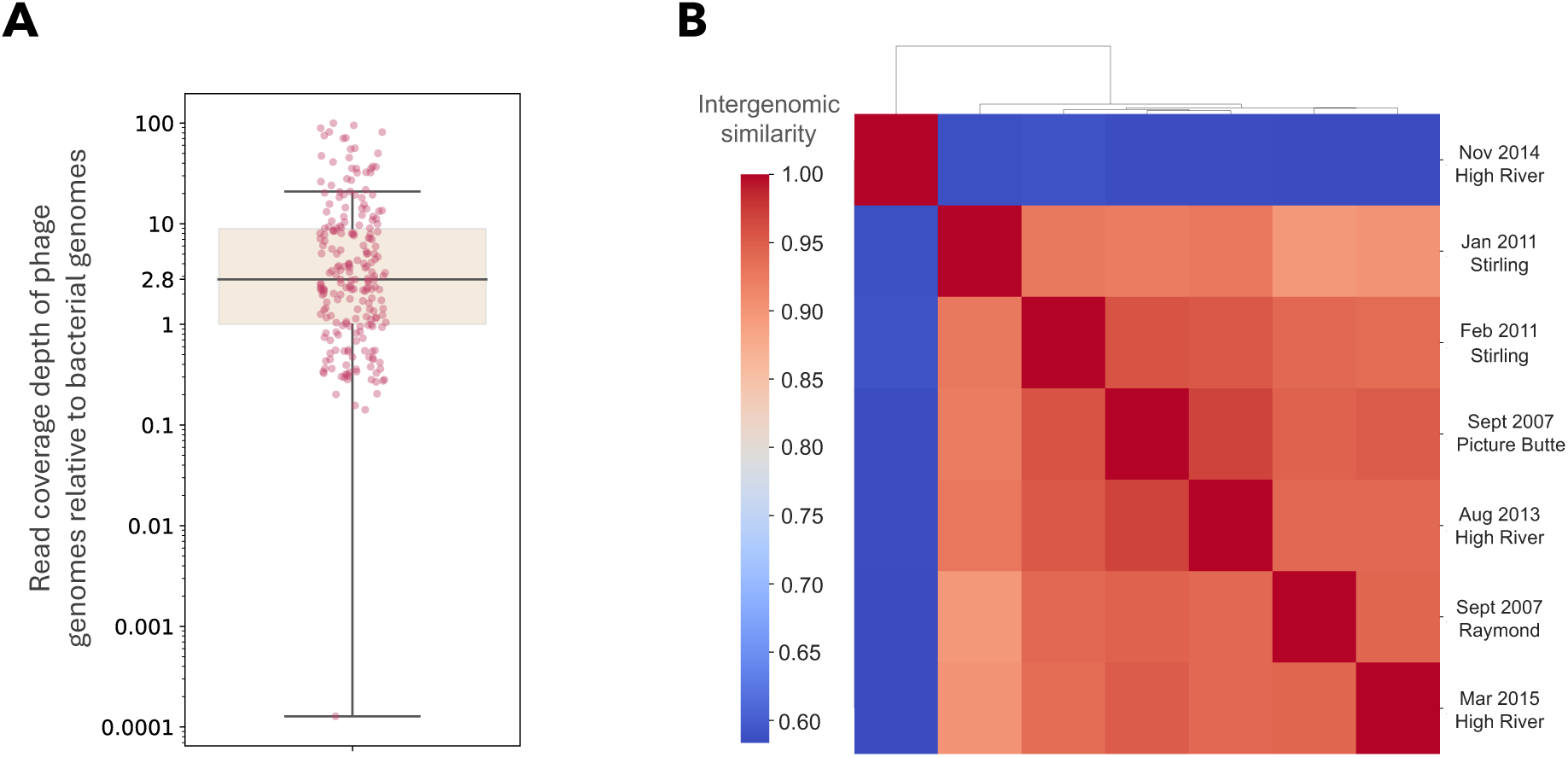
**A** Boxplot showing estimated persistent phage genome copies relative to the bacterial host for the 232 infected bacterial assemblies for which reads were available. Y-axis: read coverage depth of persistent phage contigs relative to the average of host contigs in the assembly. The box is drawn between the first and third quartiles, with a horizontal line at the median (2.8). Whiskers extend to 1.5 times the interquartile range. **B** Genome similarity between seven non-temperate phages found in Shiga toxin-producing *E. coli* bacterial assemblies published as part of a single sequencing project. Genomes are ordered and colored by their pairwise VIRIDIC intergenomic similarity, with dendrogram clustered using the nearest point algorithm. The isolation year and location of the bacterial isolates containing persistent phages is shown to the right.

We also looked at whether any labs had published multiple bacterial genomes with highly similar persistent phages, as this could be indicative of a systematic lab contamination from the same source. Excluding governmental public health projects, the NCBI BioProject with the most persistent phages was PRJNA870153, in which the authors sequenced 125 Shiga toxin-producing *E. coli* strains from cows in Alberta, Canada^31^, seven of which contained large (>350 kbp) non-temperate phages. These strains were isolated in different towns between 2007-2015 (Figure 2B), stored at -80 °C, and later thawed, cultured, and sequenced^31^. We calculated the VIRIDIC intergenomic similarities between the phages, a measure of whole-genome nucleotide similarity^32^. Only two pairs of phages were of the same respective species (>95% similarity) and none were more than 97% similar. This strongly suggesting that the viral genomes within bacterial assemblies did not stem from a lab contamination but rather from diverse environmental phages that were within the bacterial isolates.

Finally, we were interested in whether T1-like phages were present in the 373 non-temperate phage contigs. Among all phage lab contaminants, Escherichia phage T1 is perhaps the most infamous. Easily aerosolized and desiccation-resistant, T1 is known to be very difficult to eradicate in infected labs and can easily contaminate cultures^33^. However, there was no widespread T1-like contamination in our dataset, with only 2/373 non-temperate phages clustered in the same genus-level cluster as T1. From these evaluations, we found no evidence that the 373 non-temperate phages within bacterial assemblies resulted from lab contamination, with multiple lines of evidence supporting their interpretation as persistent phage infections.

### Serial propagation of infected bacterial isolates experimentally demonstrates persistent infection

Having established that the 373 persistent phages are non-temperate and likely not lab contaminates, we next experimentally tested how stable these persistent infections were. On agar plates, it is possible that an uninfected cell forms a colony that grows outwards on the plate, encountering an extracellular virion that starts replicating after the colony is already visible. If the colony is re-streaked, there may be a repeat encounter between an uninfected cell and an extracellular virion. Although this is a type of persistent infection, we wished to test whether persistent phages could co-exist in a more stable fashion. In brief, we set up a protocol where infected bacterial isolates were first inoculated from freezer stocks in liquid media followed by serial dilution onto agar plates. Colonies were then re-streaked three times each. The initial serial dilution ensured that growing colonies were very unlikely to encounter extracellular phages, and thus the persistent phage should only be present in the re-streaks if the initial colony-forming cell was infected with phage.

We selected the seven infected Shiga toxin-producing *E. coli* isolates from Alberta for this experiment (Fig. 2B). These bacteria were good experimental candidates for testing as they are infected with similar phages, were isolated and sequenced with the same protocol, and originated from the same culture collection. Applying the above protocol to all seven isolates in triplicate, we cultured 21 single colonies after dilution and three re-streaks. After sequencing, we found that one of the triplicates from isolate GCA_036253035.1 (Sept 2007, Picture Butte in Figure 2B) had 100% coverage on its persistent phage contig (JAYLNQ010000003.1). However, this was the only sample with read coverage on the persistent phage contig; only 1/21 colonies still retained their original persistent phage infection. This is a frequency far below what we might expect from a classical temperate phage but experimentally demonstrates that the persistent phages may sometimes coexist with their hosts over many generations despite rigorous attempts to cure the host.

### Persistent phages are genetically diverse and enriched in clusters of jumbo phages

Having established that the 373 persistent phages are non-temperate and likely do not represent lab contamination, we wondered whether persistent phage infections might be linked to specific host bacterial genotypes. Analyzing the taxonomy of the infected bacterial hosts, we found that 369/370 (99.7%) of the assemblies were *E. coli*, very similar to the proportion of *E. coli* in GenBank *Escherichia* genomes (99.5%). The remaining assembly was an *E. albertii* strain. Focusing on the *E. coli* strains, we constructed a core-genome phylogenetic tree of all complete *E. coli* RefSeq assemblies interspersed with the 369 infected *E. coli* assemblies (Figure 3A). Infected assemblies were distributed broadly throughout the phylogenetic tree, showing that persistent infections are not limited to certain clades of *E. coli*.

**Figure 3.**
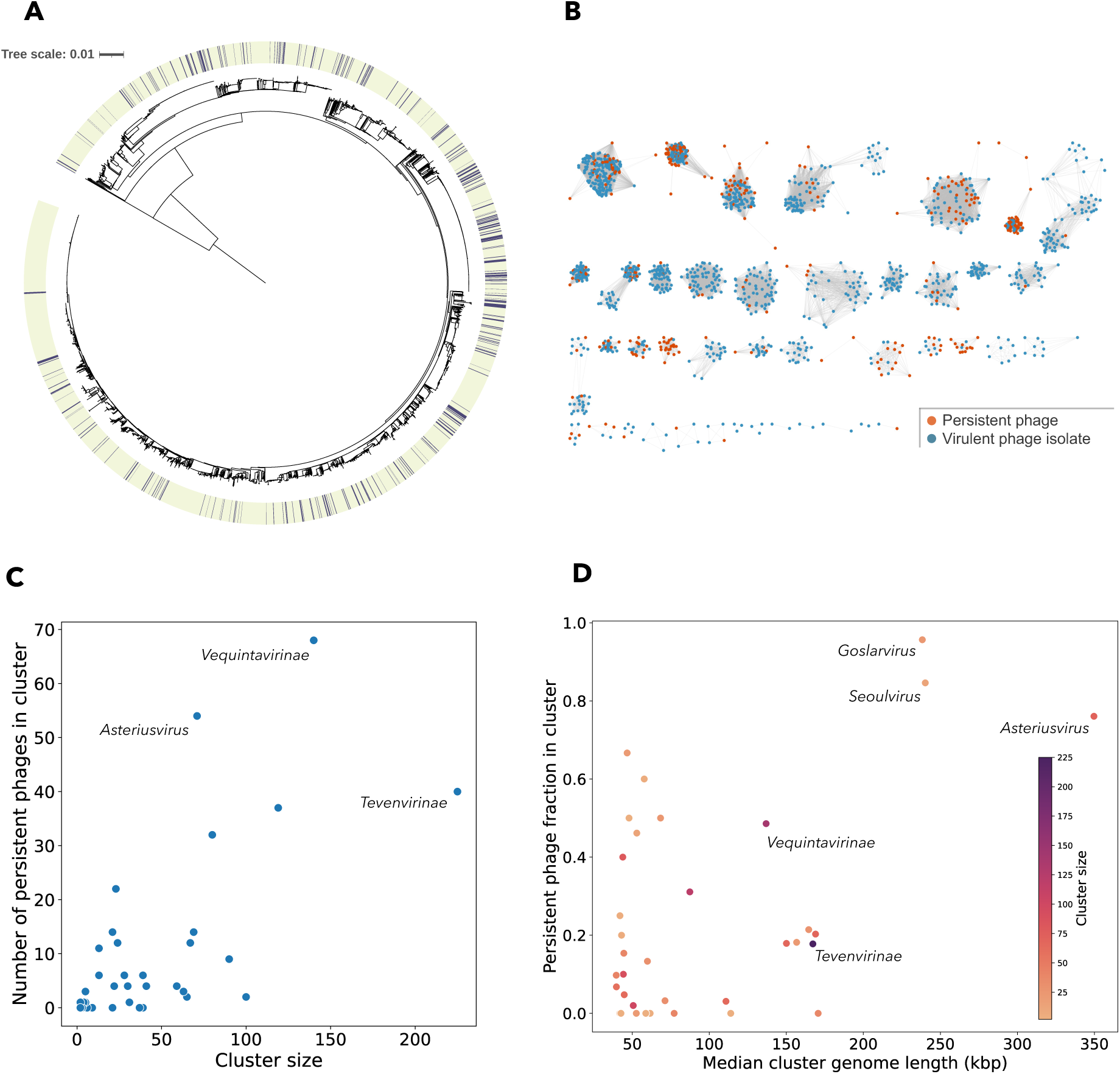
**A** Core-genome phylogenetic tree of all 3,352 closed, complete *E. coli* RefSeq genomes interspersed with the 369 *E. coli* GenBank assemblies infected with persistent phages. Rooted with the single *E. albertii* assembly with a persistent phage. **B** Protein-sharing network of virulent *Escherichia* phages isolates (blue) clustered with the non-temperate, persistent phages identified in *Escherichia* bacterial genome assemblies (orange). **C** Scatterplot showing the number of persistent phages in each of the 37 protein-sharing network clusters against the total number of phages in each cluster (cluster size). **D** Scatterplot of the median phage genome length in each cluster against the fraction of persistent phages in each cluster. Datapoints are colored by the cluster size.

Next, we investigated the genetic diversity of the persistent phages themselves. To investigate the taxonomic distribution and relative abundance of the 373 persistent phages, we analysed their relation to 1,199 virulent *Escherichia* phage isolates in a vCONTACT2 protein-sharing network (Figure 3B). The 373 phages were distributed across 152 of the 393 genus-level clusters identified by vCONTACT2, spanning a substantial fraction of the genomic diversity of virulent *Escherichia* phages. However, with an average of four genomes each, these genus-level clusters were mostly small, making it difficult to identify overrepresented clusters and further underlining the genetic diversity of the persistent phages. Using Leiden clustering to obtain larger families of phages, we defined 37 higher-level communities within the network, 27 of which contained persistent phages (Supplementary table S2). We also assigned a consensus taxonomic label for each cluster based on the ICTV taxonomy of the phage isolates. In Fig 3C, we show the number of persistent phages per cluster against the total cluster size. The cluster with the highest number of persistent phages (N=68) was the Myoviridae sub-family *Vequintavirinae,* followed by a cluster of *Asteriusvirus* (N=54) and a *Tevenvirinae* cluster containing the model phage T4 (N=40).

We also calculated the fraction of persistent phages in each cluster (persistent phage fraction). Statistically, we found that persistent phages were non-randomly distributed throughout the network clusters (permutation test, p<10^-5^), with persistent phage fractions ranging from 0 (ten clusters) to 0.96 (one cluster). Interestingly, the three clusters with highest persistent phage fraction (0.96, 0.85, and 0.76) were also the clusters with the largest genome sizes (Figure 3D). Containing the genera *Goslarvirus*, *Seoulvirus*, and *Asteriusvirus* respectively, these three were the only clusters to contain jumbo phages with genomes > 200 kbp. On the other side of the spectrum, 10 clusters contained no persistent phages, the largest of which was the sub-family *Gordonclarkvirinae* (N=39). The well-known T-phages were also in under-represented clusters, such as the *Drexlerviridae* cluster of T1 (persistent fraction 0.02), *Markadamsvirinae* cluster of T5 (persistent fraction 0.03), and the *Tevenvirinae* cluster of T4 (persistent fraction 0.18).

### Persistent jumbo phages are prevalent in bacterial taxa beyond *Escherichia*

The jumbo phage clusters containing *Asteriusvirus*, *Goslarvirus*, and *Seoulvirus* were particularly interesting as they had the highest persistent phage fractions. We therefore wondered if these non-temperate phages persistently infected other bacterial taxa beyond *Escherichia.* To examine this, we constructed core genomes for the Asterius, Goslar, and Seoul clusters and searched for similar proteins in NCBI’s non-redundant protein database. Identifying both phage genomes and contigs from bacterial assemblies enriched in these jumbo-phage proteins, we found in total 107 Asterius-like, 60 Goslar-like, and 299 Seoul-like phages (Supplementary table S3). For each of the three clusters, we calculated weighted Gene Repertoire Relatedness (wGRR) between all pairs of phages (Figure 4A-C). This is a pairwise measure of protein content similarity and therefore more broad-range than nucleotide identity, ranging from 0 (no homologous protein pairs) to 1 (all proteins in one genome have identical homologs in the other)^30^.

**Fig. 4.**
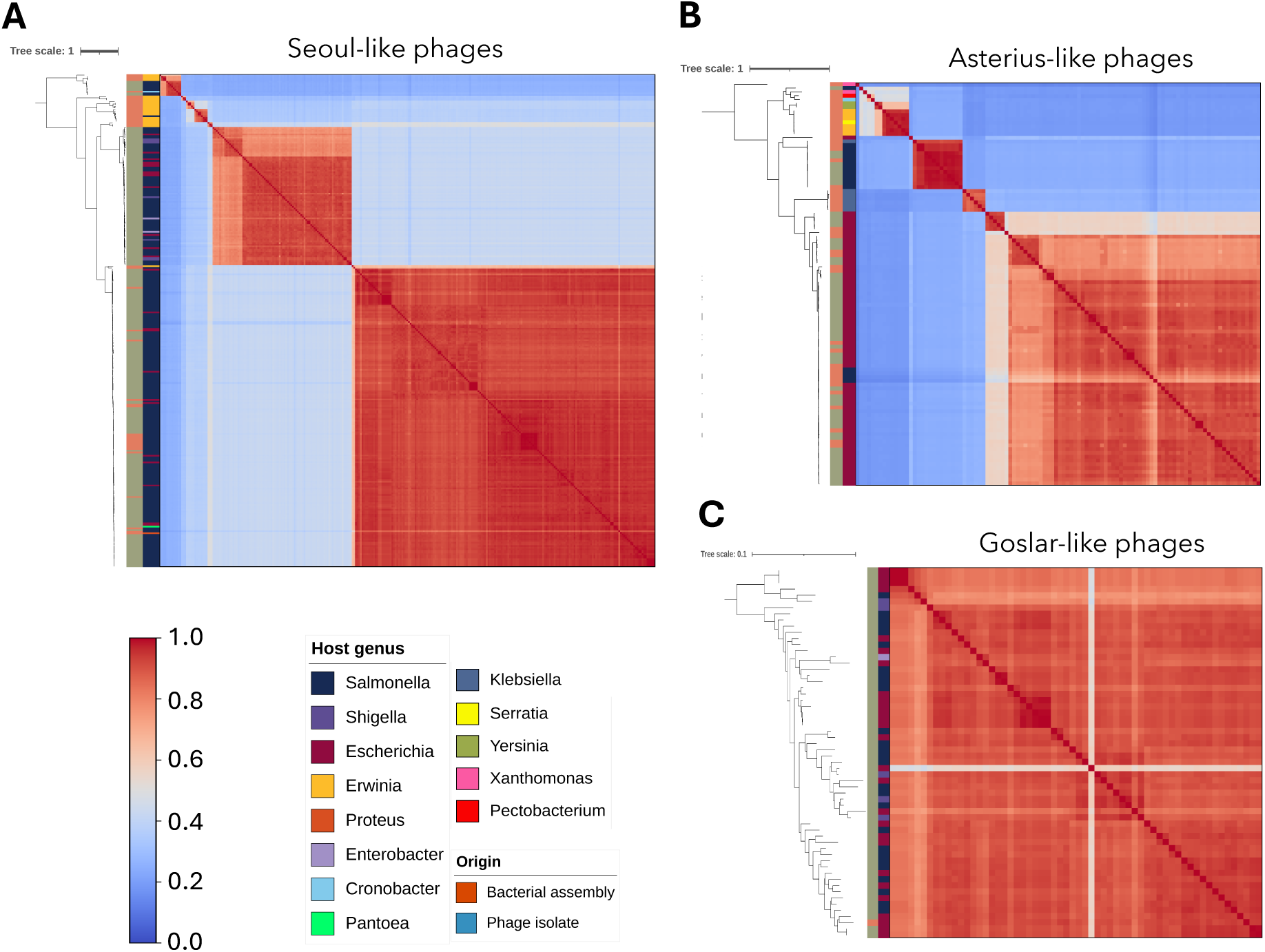
Genomes from phage isolates and bacterial assemblies in jumbo-phage clusters **A** Seoul-like phages, **B** Asterius-like phages and **C** Goslar-like phages. Phylogenetic trees were made using core-genome alignments for each cluster. Heatmaps show the weighted Gene Repertoire-Relatedness between all phages in each cluster and are ordered according to the phylogenetic trees. Color-coded columns show whether the phage genome originated from a phage isolate or bacterial assembly (1^st^), and the phage host genus (2^nd^).

The in-depth searches greatly expanded each of the clusters. The largest cluster, the 299 Seoul-like phages, was genetically diverse (wGRR pairs < 0.3) and infected bacterial hosts of families *Enterobacteriaceae* and *Erwiniaceae* (Figure 4A). Although most hosts (238) were *Salmonella*, eight genera of host bacteria were represented. In total, 84% of the Seoul-like phage cluster were persistent phages, with the persistent phage fraction almost unchanged from the initial search (85%). Meanwhile, the 107 Asterius-like phages (Figure 4B) infected bacterial hosts from nine bacterial genera of the families *Enterobacteriaceae, Erwiniaceae,* and *Xanthomonadaceae.* Asterius-like phages also had a lower persistent phage fraction (57%) than the Seoul-like phages and were more genetically diverse, containing phages with wGRR pairs < 0.2. In contrast, the Goslar-like phages (Figure 4C) are less diverse (all wGRR > 0.7), and were limited to four genera of host bacteria in *Enterobacteriaceae.* Interestingly, the Goslar-like phages have only once been sequenced as a phage isolate, with this cluster containing only a single phage genome (*Escherichia* phage Goslar) together with 59 persistent phages derived from bacterial assemblies.

The genera *Goslarvirus* and *Seoulvirus* both reside within the family *Chimalliviridae* of nucleus-forming phages and are distantly related to Pseudomonas phage phiKZ^34^, while the genus *Asteriusvirus* is unrelated and does not have an assigned taxonomic family. To confirm that members of the expanded Goslar-like and Seoul-like clusters all encode the phage nuclear protein chimallin, we searched for hits to the experimentally confirmed chimallin from Goslarvirus Goslar (YP_009820873) and Seoulvirus SPN3US (YP_009153538)^35^. Chimallin was very well conserved in these clusters. All 299 Seoul-like phages encoded a protein with > 70% similarity to SPN3US chimallin, while 59/60 Goslar-like phages had hits with > 90% similarity to Goslar chimallin. This suggests that the Seoul-like and Goslar-like phages have the potential to form phage nucleii during infection.

We also found interesting indications of long-term phage-host coexistence within the Seoul-like phages. Although the Seoul-like cluster contained very diverse phages (pairs with < 0.3 wGRR), there were also highly similar pairs of phages. Notably, 35 pairs of phage assemblies had wGRR = 1 (not necessarily identical at the nucleotide level). If we were to find the same persistent phage genome in multiple isolates of a bacterial strain, it would suggest a long-term persistent infection outside the lab. To investigate this, we contacted the UK Gastrointestinal Bacteria Reference Unit for metadata on five such pairs of very similar persistent phages, each derived from clinical *Salmonella* isolates in their collection. Although four of these isolate pairs were isolated from the same respective patients, one pair of *S. enterica* Braenderup strains (GenBank accessions ABSICT000000000 and ABSGIM000000000) were confirmed to be isolated from different patients in 2021 and 2024 (personal communication). Interestingly, both the bacterial hosts and the persistent phages in these two isolates were 100% identical; we performed variant-calling using sequencing reads from the two assemblies but found no variants (SNPs or Indels). This result suggests that the Seoul-like phage may have co-existed with its bacterial host over a period of three years, and that phage infections may persist long-term in environments outside the lab.

## Discussion

Although there are many reports of phages with lifestyles that do not fall into the traditional virulent/temperate dichotomy^19,20^, we have little idea of how common they are. Here we conduct the first systematic survey of such phages by searching bacterial assemblies for known virulent phages. We also experimentally demonstrated that a non-temperate jumbo phage can persist long-term with an infected bacterial host. These results expand the notion that a clonal bacterial culture is not a static, single entity, but rather an ecosystem of genetic elements. In addition to known mobile genetic elements such as inducible temperate phages, plasmids, and IS elements, we here show that many “pure” bacterial cultures contain undetected virulent phage infections.

Although our results show that certain groups of phages are more prone to persistent infections than others, it is unclear what mechanisms underlie the persistent phage phenotype. Here we suggest three possible mechanisms: **phase variation**, **asymmetric phage inheritance**, and **lysis inhibition**.

### Phase variation

Phage-bacterial coexistence could result from phase variation in bacterial phage receptors leading to a mixed bacterial population of susceptible and resistant cells (Fig. 5A)^36^. Documented examples of phase variation in potential phage receptors include reversible fimbriae expression in *E. coli*^36^ and O-antigen chain length variation in *S. enterica,* which has been shown to influence phage resistance^37^. Phase variation is not limited to receptor genes; a restriction modification system in *Campylobacter jejuni* was shown to be phase-variable, impacting phage resistance^38^. Similarly, phase variation of capsular polysaccharides in *Bacteroides intestinalis* is associated with long-term coexistence with the phage crAss001^17^. Through the continuous generation of heterogeneous bacterial populations containing a mix of sensitive and resistant bacteria, phase-variation could result in long-term co-existence of phage and bacteria. A mechanism where phage replication is limited to a subset of the bacterial population would also explain the moderate phage-host DNA ratios we observed.

**Fig. 5.**
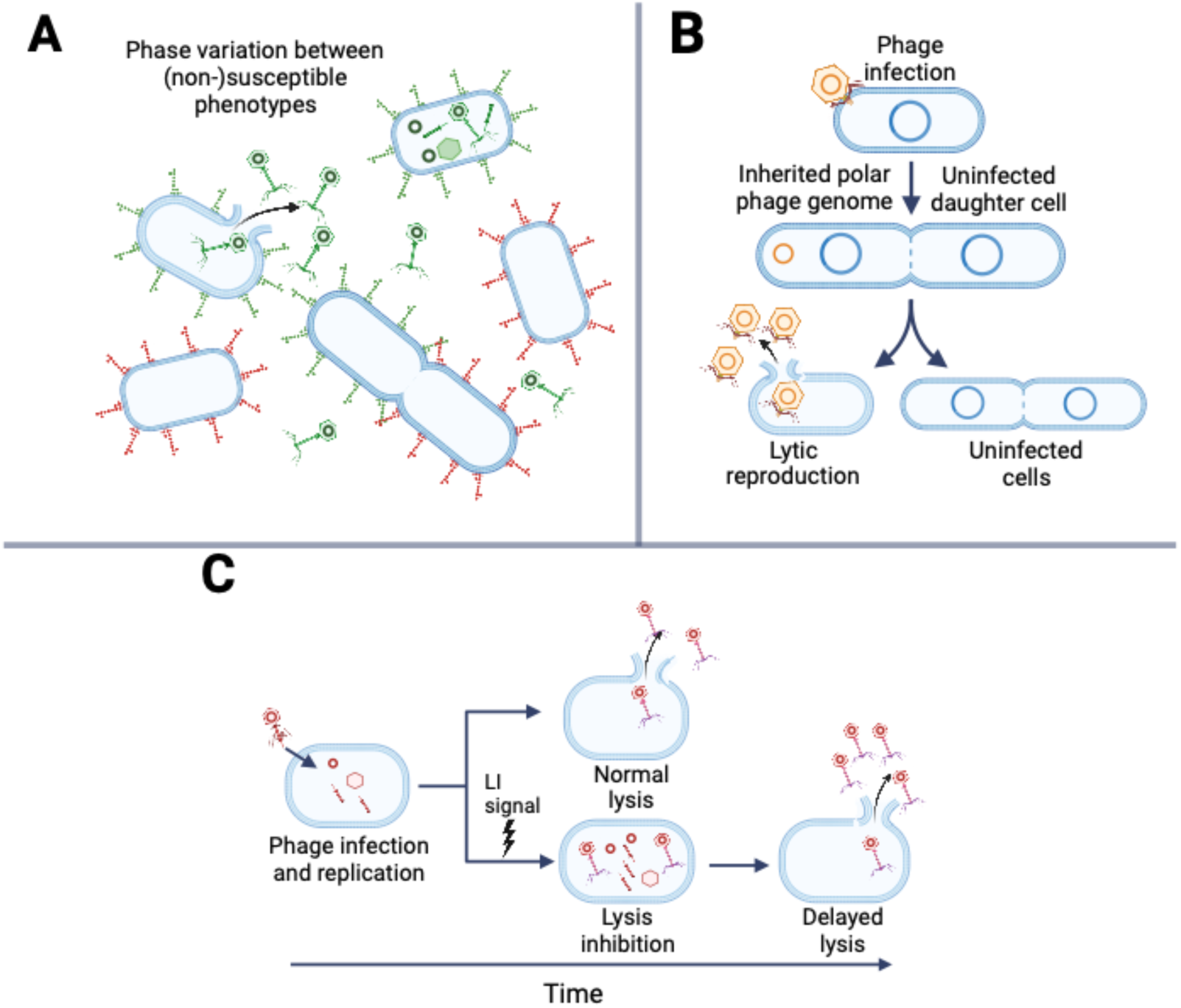
Proposed mechanisms for persistent infection. **A** Phage variation **B** Asymmetric phage inheritance **C** Lysis inhibition. Created with biorender.com

### Slow growth and asymmetric phage inheritance

Bacterial populations with a mix of susceptible and resistant cells may also be generated through other means. If phage replication is slow and infected bacterial cells continue to divide, phage DNA may be asymmetrically inherited between mother and daughter cells, resulting in a mixed population of infected and uninfected cells (Fig. 5B). Repeated over many generations, this may result in a persistent infection. This has been observed in Salmonella phage P22, a temperate phage which can delay its lysis/lysogeny decision by remaining as an episomal genome through cell division with only one of the daughter cells inheriting the phage genome. The uninfected daughter cell is temporarily immune to subsequent phage infection due to inheritance of a phage-encoded repressor protein from the mother cell, sustaining an uninfected bacterial subpopulation^14,39^. Once this repressor is sufficiently diluted, the cell is re-sensitized to infection and the cycle can repeat itself. Some jumbo phages have been shown to replicate slowly, which could contribute to the overrepresentation of large phages in persistent infections. For example, the *Pseudomonas* jumbo phage PA5oct was shown to persist in some bacterial cultures where phage DNA was replicating slower than that of their bacterial hosts^40^.

### Lysis inhibition

Lysis inhibition as described in *Escherichia* phage T4^9^, is a process where intracellular phages begin replication, but delay host cell lysis in response to signals such as high MOI (Fig. 5C). The jumbo phiKZ-like phages may also be capable of lysis inhibition, with reports that *P. aeruginosa* colonies infected with jumbo phage phiKZ at high MOI can continue to grow for several days, turning opalescent in color, before eventually lysing and releasing high numbers of phage progeny^12,13^. Phage phiKZ is part of a family of very large viruses (Chimalliviridae)^34^ that form protective nuclear compartments constructed from chimallin during infection, shielding the replicating phage from bacterial defense systems. Here, we confirmed that the Seoul- and Goslar-like persistent phages also encode the major nuclear protein chimallin. Although the moderate phage-host DNA ratios we observed mostly rule out scenarios where all bacterial cells are packed with full lysis-inhibited phages as in fully opalescent phiKZ-infected colonies, this could be explained if DNA was isolated from the infected cultures at an earlier stage in the phage lysis inhibition. Although the phage nucleus has not been linked to lysis inhibition or persistent infection, the overrepresentation of chimallin-encoding phages in our dataset is intriguing.

Regardless of mechanism, persistent infection may well be a more common phenomenon than our results suggest. Given our search strategy’s reliance on similarity to sequenced phage isolates, we certainly did not detect all persistent infections in *Escherichia* assemblies, as phages dissimilar to known isolates will have gone undetected. Even in a well-studied genus like *Escherichia* there are doubtlessly many undiscovered phage clades, especially those that are difficult to isolate with plaque assays such as persistent phages. Combining the results from the initial search in *Escherichia* with the subsequent targeted search for the three jumbo phage clusters, we identified 658 high-confidence virulent phages in bacterial assemblies. Our results from the targeted search for *Asterius*-, *Seoul-*, and Goslar-like viruses also demonstrate that persistent infections occur outside of *Escherichia*, although they may be more difficult to detect in taxa with fewer sequenced phage isolates upon which to base the search.

These results may be relevant to the selection of phages in therapeutic applications. Temperate phages are generally avoided in therapy in part due to the presumed lack of efficient bactericidal effect^41^, and it does seem likely that phages from clusters with very high persistent phage fractions may be less effective at eliminating their hosts. In recent years, jumbo phages have attracted a lot of attention as potential therapeutic tools due to their often-times broad host ranges^42–44^, including phages within the Seoul-like^45^ and Asterius-like clusters^46^. Although some such phages are capable of bactericidal effect under certain conditions, the possibility that they may result in persistent, non-lethal phage infection might have to be considered before using them as therapeutic agents.

This work joins other studies in suggesting that classification of phages as either virulent or temperate may be too simplistic^20,47^. While some virulent *Escherichia* phage clusters were dominated by either phage isolates (i.e. *Gordonclarkvirinae)* or persistent phages (i.e. Goslar-like viruses), most clusters were somewhere in-between, such as the *Tequatrovirinae* cluster containing the archetypal virulent phage T4 (persistent fraction 0.18). Although persistent infections clearly act through different mechanisms than true temperate phages (they do not form prophages), a persistent infection does resemble a temperate phage infection in that phage-host coexistence results in a reduced host burden (lower virulence). Asymmetric division and lysis inhibition can even protect infected host cells from subsequent phage infection like temperate phages do, potentially benefiting the host. Given the existence of phage clusters with both virulent and persistent phages, we suggest that persistent infection is a variable, intermediate phenotype dependent on host and environmental conditions, similar to how the induction rates of true temperate phages vary in response to cellular and external signals^7,48^. With this interpretation, we propose that phage-host interactions, both for temperate and non-temperate phages, exist on a gradient of virulence, with rapid, lytically replicating virulent phages on one side, and stable, non-inducing prophages on the other.

## Methods

### Search for *Escherichia* assembly contigs resembling virulent phages

We first set out to identify contigs in *Escherichia* isolate genome assemblies that resemble virulent phage genomes. To do so, we used two main GenBank datasets: complete phage genomes (27,022 genomes, retrieved March 2024), and *Escherichia* bacterial genome assemblies (267,902 annotated assemblies, retrieved May 2024). In brief, we searched for contigs in the bacterial assemblies that resembled virulent GenBank phage genomes (Figure 1).

In more detail, we classified the lifestyle of the GenBank phage genomes using BACPHLIP v0.9.6^24^ as either virulent or temperate (virulent if score > 0.5), with 18,489 phage genomes labelled virulent. Using Prodigal v2.6.3^49^ we predicted 1,766,709 proteins from these virulent phages. We then clustered the proteins from virulent phages with all 1,148,448,116 bacterial proteins from the *Escherichia* assemblies using MMseqs2 linclust v14.7e284^50^ at 80% identity and coverage, yielding 894,667 protein clusters. For each bacterial contig, we counted the number of encoded proteins clustered with proteins from virulent phages. To select contigs enriched in virulent protein clusters, we retained contigs with > 25 proteins and where > 70% of their proteins were in virulent protein clusters (28,819 contigs).

We then filtered the 28,819 contigs to remove those with proteins that resembled temperate phages more than they resembled virulent phages. First, we re-clustered proteins from the 28,819 contigs with all GenBank phage proteins using MMseqs2 linclust as before. For each bacterial contig, we identified the most similar phage genome, quantified by the number of shared MMseqs2 protein clusters. We discarded the contigs most similar to temperate phage genomes, keeping only the 5,234 contigs from 5,030 bacterial assemblies whose closest phage isolate was virulent (predicted by BACPHLIP).

We also screened the remaining contigs for similarity to 3,514 complete (gapless chromosomes and plasmids) RefSeq *Escherichia* assemblies (retrieved May 2024). As virulent phages do not integrate into bacterial chromosomes, we assumed virulent phages should not be found in complete assemblies. Similarity to complete assemblies was therefore used as an indication that the bacterial contig may resemble temperate phages integrated in the complete assemblies. Using sourmash v4.8.8^25^, we checked whether the 5,234 contigs were contained within any of the complete RefSeq assemblies using parameters kmer length=31 and scaled=1000. 4,726 contigs with containment > 0.15 were labelled putative temperate phage contigs, while the remaining 508 contigs were labelled putative virulent phage contigs.

### Identification of 373 high-quality virulent phages in *Escherichia* assemblies

Next, we used vCONTACT2 v0.11.3^51^ to cluster the 5,234 labelled contigs together with 1,331 GenBank *Escherichia* phages (all phages with > 25 proteins and “Escherichia” in their name), clustering into 462 viral clusters (VCs). 142 bacterial assemblies contributed multiple contigs to this dataset (346 contigs in total). Although we expected virulent phage contigs from the same bacterial assembly to represent the fragmented genome of a single virulent phage, only 1/142 of these assemblies had contigs that clustered in the same VC. To improve this, we trimmed the similarity network generated by vCONTACT2, normalizing similarity between 0-1 and retaining similarities > 0.1, and used this as input for Leiden clustering^27^ with resolution parameter 1 (implemented with igraph v1.2.11^52^ and leidenalg v0.10.0). Using Leiden clusters, 75/142 assemblies with multiple contigs had their contigs clustered within the same VC. We binned all contigs from the same bacterial assembly that clustered together into multi-contig phage genomes, resulting in 5,106 combined phage genomes from bacterial assemblies.

We then conducted a subsequent round of vCONTACT2, clustering the 5,106 binned phage contigs from *Escherichia* bacterial assemblies with the 1,331 *Escherichia* phage isolate genomes. Here, vCONTACT2 clustered 5290/6437 phages into 443 VCs. Generating Leiden clusters (LCs) as before clustered 6361 phages into 100 LCs which were used for downstream analysis. The 64 contigs from assemblies that formed cluster singletons were discarded and not considered further.

We calculated the fraction of virulent phage genomes in each LC (both from phage isolates and bacterial assemblies) using the sourmash-derived labels for the binned phage contigs, and the BACPHLIP-derived labels for the phage isolates. Only eight LCs contained mixed virulent and temperate phage labels, with all other clusters unanimously virulent or temperate. All unanimously temperate clusters were discarded. 6/8 mixed clusters were discarded, as although they contained BACPHLIP-labelled virulent phage isolates, the clusters also contained contigs from bacterial assemblies with sourmash hits to complete bacterial chromosomes. The two remaining mixed clusters were retained. One (cluster 52) was retained despite high sourmash similarity to a single RefSeq plasmid (NZ_CP067310.1) as annotation did not reveal any plasmid-specific genes and all BACPHLIP predictions in the cluster were virulent. The second (cluster 26) was also retained; although 19/55 BACPHLIP predictions in cluster 26 were temperate, there was no similarity to complete RefSeq assemblies, no obvious temperate genes and no reports in literature of the two ICTV genera within the cluster (*Phapecoctavirus* and *Justusliebigvirus*) being temperate^53–55^.

Finally, CheckM v1.1.6^56^ was used to evaluate contamination in the assemblies. Four assemblies with > 5% contamination were removed. After this, we had 493 contigs from 370 *Escherichia* bacterial assemblies binned into 373 non-temperate, virulent phage genomes.

### Search for plasmid and pseudolysogeny genes

We searched for plasmid functions in the 373 persistant phage genomes using established HMM models for nine partition systems and 38 replication proteins^30^. To select for hits that covered most of the gene and not a single domain, we filtered hits with E-value < 5e-5 and model coverage >= 50%.

### Calculation of persistent phage relative read coverage depth

232 of the GenBank *Escherichia* bacterial assemblies containing non-temperate, persistent phages had read data available. We used BWA-MEM 0.7.18^57^ to map the reads back to their respective assemblies and samtools 1.18^58^ to calculate the average sequencing depth of each contig. The weighted sequencing depth average was calculated for contigs identified as virulent phages, in addition to a weighted average for all bacterial contigs. From these two numbers, the read coverage depth relative to the bacterial genome was calculated for each persistent phage.

### Stability test of *E. coli* isolates infected with persistent phages

The stability of persistent Asterius-like phages was tested for seven *E. coli* isolates in the following manner. Each isolate was inoculated from -80 °C stocks into 4 mL EC broth and incubated at 37 °C for 24 h. The seven cultures were then serially diluted, plated onto EC agar plates and incubated at 37 °C for 24 h. From agar plates with at most 10 colonies, three well-separated colonies were picked from each plate and re-streaked onto separate agar plates (triplicate colonies from each of the seven isolate dilutions). The 21 re-streaked plates (re-streak 1) were incubated at 37 °C for 24 h. One well-separated colony from each plate was then re-streaked onto new plates (re-streak 2) and incubated as before. This was repeated once more (re-streak 3). From the final round of 21 plates, one well-separated colony was picked and inoculated in EC broth, incubating at 37 °C for 24 hours. We then isolated DNA from each of the 21 cultures for ONT long-read sequencing. Samples were barcoded with the Rapid Barcoding Kit 24 V14 (ONT, UK) and sequenced on a PromethION R10.4.1 flow cell (ONT, UK). Reads were basecalled and demultiplexed using dorado 0.8.2 with the dna_r10.4.1_e8.2_400bps_sup@ v5.0.0 model for simplex reads and the dna_r10.4.1_e8.2_5khz_stereo@v1.3 model for duplex reads. Reads were then mapped to the persistent phage contigs in the original isolate assemblies using minimap2 2.24-r1122^59^. Read coverage statistics were then calculated using SAMtools^58^.

### *E. coli* phylogenetic tree

Next, we constructed a phylogenetic tree of the 369 *E. coli* isolate assemblies containing persistent phages together with all 3,352 complete *E. coli* RefSeq assemblies. The tree was rooted with the *E. albertii* assembly ASM2756809v1 which also contained a persistent phage. Using PanaCoTA v1.4.1^60^ we clustered all proteins at 80% identity and coverage. Constructing and aligning a core genome (99.9% presence) of 443 genes, we used FastTree v2.1.11^61^ with the default Jukes-Cantor + CAT model to generate a phylogenetic tree. RefSeq assembly GCF_021307385.1 was removed as it looks to be misclassified. iTOL v6^62^ was used to visualize the tree.

### HMM search for additional phages

We searched for additional phages related to the three *Escherichia* clusters of Asterius-like viruses, Goslar-like viruses and Seoul-like phages. First, we ran CheckV v1.0.3^28^ on the persistent phage genomes, removing medium- and low-quality contigs (potentially incomplete phage genomes). Next, we used PanaCoTA to construct core genomes for each cluster using both the phage isolate genomes and the high-quality/complete persistent phages. Proteins were clustered at 80% coverage and 25% identity, with clusters defined as core if they were present in >= 90% of genomes. The Asterius core genome consisted of 334 genes, the Goslar core genome of 214 genes, and the Seoul core genome of 219 genes. MAFFT v7.526^63^ was used to align core genes and HMMER v3.3.2^29^ was used to build hidden Markov models for each core gene and search for hits in NCBI’s non-redundant protein database (retrieved July 2024). Hits were filtered at >80% coverage and >250 score. We retrieved the GenBank nucleotide assemblies (both bacterial and phage genomes) encoding each protein hit, counting the number of core gene hits per assembly. 819 assemblies with at least five core gene hits to one of the clusters (Asterius, Goslar, or Seoul) were downloaded for further analysis.

We then built BLAST protein databases containing all proteins in the Asterius, Goslar, and Seoul clusters respectively. Next, we used BLASTP v2.14.1+^64^ to search all proteins in the 819 bacterial assemblies against their relevant Asterius/Goslar/Seoul protein databases. Protein hits were assessed using a cutoff of 80% coverage and 25% identity. Bacterial contigs with > 10 protein hits were then filtered by their percentage of total protein hits, with thresholds determined after manual inspection (40% hits for Asterius, 80% for Goslar, and 50% for Seoul). Phage contigs from the same bacterial assembly were binned into single phage genomes. Asterius phage genomes with combined length < 300kbp, and Goslar and Seoul genomes < 200kbp were removed to retain only complete or nearly complete assemblies. This left 107 Asterius genomes, 60 Goslar genomes, and 299 Seoul genomes for further analysis.

Core genomes for the expanded jumbo phage clusters were again constructed using PanaCoTA (80% coverage, 25% identity and present in >=95% genomes). After MAFFT protein alignment, IQ-TREE -v2.2.5 with automatic model selection (Asterius: Q.yeast+F+I+G4, Goslar: JTT+F+I+G4, and Seoul: Q.insect+F+I+G4) was used to generate phylogenetic trees for each cluster. Trees were visualized with ITOL v6. Weighted Gene Repertoire Relatedness (wGRR)^30^ was also calculated between all genome pairs in each cluster and visualized in a heatmap.

### Variant calling between two *S. enterica* assemblies infected with Seoul-like viruses

Two *S. enterica* isolates confirmed to be isolated from separate patients (GenBank accessions ABSICT000000000 and ABSGIM000000000) were found to contain very similar Seoul-like phages. We checked the similarity of both bacterial host and associated persistent phages by downloading the SRA read datasets associated with one of the assemblies (ABSGIM000000000) and variant-calling against the other assembly (ABSICT000000000) using snippy 4.6.0^65^.

## Acknowledgements

This work was funded by EMBO Scientific Exchange Grant 10709, Novo Nordisk Foundation grant NNF19SA0059348, RESISTE ANR-20-CE35-014, INCEPTION project (PIA/ANR-16-CONV-0005), Laboratoire d’Excellence IBEID Integrative Biology of Emerging Infectious Diseases [ANR-10-LABX-62-IBEID]. This work also used the computational and storage services (TARS cluster) provided by the IT department at Institut Pasteur, Paris.

We also thank Dr Marie Chattaway and Dr Claire Jenkins for their help with strain metadata.

